# Modeling the effects of *Aedes aegypti*’s larval environment on adult body mass at emergence

**DOI:** 10.1101/2021.05.24.445402

**Authors:** Melody Walker, Karthikeyan Chandrasegaran, Clément Vinauger, Michael A Robert, Lauren M Childs

## Abstract

Mosquitoes vector harmful pathogens that infect millions of people every year, and developing approaches to effectively control mosquitoes is a topic of great interest. However, the success of many control measures is highly dependent upon ecological, physiological, and life history traits of the mosquito species. The behavior of mosquitoes and their potential to vector pathogens can also be impacted by these traits. One trait of interest is mosquito body mass, which depends upon many factors associated with the environment in which juvenile mosquitoes develop. Our experiments examined the impact of larval density on the body mass of *Aedes aegypti* mosquitoes, which are important vectors of dengue, Zika, yellow fever, and other pathogens. To investigate the interactions between the larval environment and mosquito body mass, we built a discrete time mathematical model that incorporates body mass, larval density, and food availability and fit the model to our experimental data. We considered three categories of model complexity informed by data, and selected the best model within each category using Akaike’s Information Criterion. We found that the larval environment is an important determinant of the body mass of mosquitoes upon emergence. Furthermore, we found that larval density has greater impact on body mass of adults at emergence than on development time, and that inclusion of density dependence in the survival of female aquatic stages in models is important. We discuss the implications of our results for the control of *Aedes* mosquitoes and on their potential to spread disease.

**Author summary:** In this work we examined how the environment in which young mosquitoes develop affects their adult body size as measured by adult body mass. Adult size has potential impacts on mosquito behavior and the ability of mosquitoes to transmit disease. We used a combination of experimental work and mathematical modeling to determine important factors affecting adult mosquito body size. In our model, we incorporated potentially interacting aspects of the mosquito life cycle and traits that affect mosquito growth as juveniles. These aspects include body mass, density of the population, and level of available resource. We compared different models to determine the one that best describes the data. As mass at emergence is linked to the success of adult mosquitoes to produce offspring and to their ability transmit pathogens, we discuss how important influences on development and survival of young mosquitoes affect mosquito control and disease spread.

## 1 Introduction

Mosquito-borne diseases pose a significant global health threat, impacting over 300 million people each year [1]. Historically, control of most mosquito-borne diseases focuses on decreasing mosquito population size [2]. Common control strategies include applications of insecticides [3], but new population reduction interventions such as those that include releases of *Wolbachia*-infected mosquitoes that induce females sterility [4] and genetically modified mosquitoes that pass on lethal genes, decrease sterility, inhibit flight, or change males to females [5–8] are being tested and implemented across the globe [4,8]. While these traditional and novel interventions directly alter the adult mosquito population, it is essential to consider the potential effects on juvenile stages as well because many important morphological processes occur in the early developmental stages. These processes can significantly influence mosquito life history characteristics and the role of mosquitoes in pathogen transmission.

While mosquitoes follow a similar life cycle, details can be species specific; in this work, we focus on the genus *Aedes.* Mosquitoes use plant-derived sugars (e.g. floral nectar, honeydew, fruits, etc.) as their main form of nutrients, but female mosquitoes require blood meals to develop eggs. Thus, only female mosquitoes bite vertebrates. Once a female acquires sufficient reserves of blood, she lays eggs on the surface of water [9]. Some species including *Aedes aegypti* and *Aedes albopictus* lay their eggs in small holes or man-made containers (e.g. tires or rain barrels), and a single egg batch typically ranges from approximately 50 to 100 eggs [10,11]. Once the eggs are sufficiently hydrated, they hatch into larvae. The larvae progress through four stages called instars, becoming larger in each stage, and ultimately molting into a pupa. Development time from hatching of an egg to emergence as an adult depends on several factors including food, density, and temperature [9,12]. During the pupal stage, the mosquitoes do not feed [13], and after a short time, pupae emerge as adult mosquitoes. The total time from hatching of an egg to adult emergence is highly variable and can range from as short as 7 days to more than 90 days [9,14].

The conditions experienced during each juvenile stage differentially affect outcomes later in the mosquito’s life, such as age and mass at emergence. In particular, mosquito body size is correlated with several adult traits such as fecundity and longevity [10,15–19]. Studies show a positive correlation with body size and fecundity, such as Briegel’s work that found that female *Aedes aegypti* with larger body sizes had two to three times as many mature eggs as smaller mosquitoes [17]. Large males, too, have been associated with greater fecundity in females who mate with them versus those who mate with small males [20,21]. In regards to survival, larger mosquitoes were shown to live longer on average [19,22–24]. In a study on competition between *Aedes aegypti* and *Aedes albopictus*, body size was found to be a significant indicator of survival regardless of the competition levels [22].

Body size also affects the behavior of adult mosquitoes. Larger mosquitoes were shown to be more successful at blood feeding [25] and more persistent in acquiring a sufficient blood meal from a particular host [26]. In contrast, small mosquitoes are less persistent on a particular host and often do not obtain a complete blood meal in a single bite. Thus, smaller mosquitoes frequently bite more times and across more individual hosts [10,26]. Additionally, large mosquitoes take larger blood meals [27]. These relationships between body size and feeding behavior can impact the level and spread of mosquito-borne disease. For example, Juliano et al. showed important relationships between body size and vector competence in *Aedes aegypti*, the primary vector of dengue [28]. In particular, field collected mosquitoes were more likely to be infected with dengue if they were larger in size [28]. Furthermore, Bara et al. observed that smaller mosquitoes showed greater dissemination rates of dengue compared to larger mosquitoes [29].

Despite the rich modeling literature on mosquito population dynamics, intricacies of the early stages of mosquito development are often ignored, and only a few modeling studies focus on mosquito body size in *Aedes* [30–32]. However, some work has been done in *Anopheles,* including a study investigating density and body size. In this work, a statistical model demonstrated the importance of larval density dependence and maternal wing length on the adult population size of the subsequent generation, and suggested that higher density might be one factor that leads to smaller mosquitoes [33]. However, this model did not include any direct interaction between larval density and body size. In another study, larval environments of *Anopheles* were found to be significantly associated with adult size, longevity, and the ability to spread malaria [34]. Using a differential equation model of *Anopheles,* Lunde et al. showed that the average adult wing length in the population was correlated with population size [35]. In this study, the wing length of newly emerged adults was a linear function of increasing larval body size, and average body size was assumed to decrease linearly with increasing larval density. Adult body size, temperature, and humidity were included as independent variables affecting mortality. Lunde et al. compared this model with five other models excluding size and found that the model including size did better than all of the other models predicting mortality rates [36]. However, using their mathematical model, they determined that size was not as influential as temperature and humidity in adult mortality. In several studies involving *Aedes*, wing length was used to estimate the number of eggs laid by a female in a metric to measure per capita intrinsic population growth rates [15,16,37,38], which was first introduced by Livdahl and Sugihara [39]. While progress has been made towards understanding the role of adult body size in mosquito population dynamics, important questions remain about how larval population dynamics impact adult body size.

In 1979, Gilpin and McClelland developed one of the first models to consider how the mass of *Aedes aegypti* larvae changes due to food resources [31]. They assumed that the increase of weight in larvae is proportional to a power of weight and a Holling type two function of food. In particular, they formulated equations that dictated the amount of food that is converted into mosquito mass. The food available in the system was reduced by this amount for each of the larvae present at the time, and the total mass increased by a proportion of the amount of food reduction. They also assumed that larvae must reach a minimum weight and persist for a minimum time before they pupate. This framework is the foundation of several models that include resource-dependent juvenile population dynamics and impacts on the body size of *Aedes* mosquitoes [40,41]. Padmanabha et. al. considered that reserves, i.e. stored energy that can be used for needs (excluding weight that is structural), were more important than weight itself [32]. They concluded there is a point in the final larval stage (L4) at which larvae commit to pupate, i.e. become a pupae. The authors used a growth model which was maximized at a threshold of current reserves, and where growth was diminished farther from the peak value. They used a logistic function to describe mass over biological time with the peak change in growth occurring during the final larval stage. They compared models with growth dependent on reserves to growth dependent on mass and showed reserves to be more important in modeling growth. In a more recent study, Aznar et al. compared the Gilpin-McClelland model with a compartment model [30]. They considered that development time of each biological stage was gamma distributed. The variance of the gamma distribution was food dependent for some stages, but food independent for others. They found a quadratic relationship between the variance and mean, where the mean development time changed with food availability. This larval model was also extended and used to study impacts of the sterile insect technique on a wild population [42].

In this work, we consider populations of *Aedes aegypti* mosquitoes, vectors of the viruses that cause dengue, Zika, chikungunya, and yellow fever, among other diseases [43]. These mosquitoes are known to lay eggs in small natural or man-made containers such as tires, buckets, and tree holes [44,45]. Given the inherent constraints on resources of container habitats, larger larval populations can experience greater competition for space, food, and other resources than smaller populations [46,47]. As a result, this often leads to morphological changes in emerging adult mosquitoes such as variation in adult body size [48]. Because body size is important to adult mosquito life history characteristics and a mosquito’s potential as a vector of pathogens, it is important to develop a better understanding of contributions to adult body size of competition and density dependence in larval populations. Herein, we utilize data collected from experiments aimed at characterizing relationships between larval environment and adult body size to develop a discrete time mathematical model that accounts for effects of density and resource availability on larval mass and, ultimately, on adult mosquito body mass. We are particularly interested in creating a model that will produce variability in sizes based on larval environments. We collected experimental data to inform structure and parameter values of multiple possible models, and we compare fit across models to determine which model is most appropriate. For any parameter not fit to data, we consider variations and sensitivity of the model to that parameter. Finally, we discuss the importance of our results for mosquito life history characteristics, pathogen transmission, and efforts to control mosquito populations.

## 2 Materials and Methods

### 2.1 Data Collection

Data was collected from a study of the mosquito species *Aedes aegypti*. We conducted two types of experiments. In the first, we tracked development through each aquatic stage under low and high larval density conditions. In the second, we tracked total length of development time and mass upon emergence under low and high larval density conditions.

We study *Aedes aegypti* mosquitoes due to their medical relevance [49]. We used the Rockefeller strain (MR-734, MR4, ATCC ®, Manassas, VA, USA) due to the extensive literature on their behavior, which makes comparisons with other studies easier [50]. Larvae were reared in densities of 26 and 78 per 300 mL of nutrient medium to simulate ‘low’ and ‘high’ intraspecific competition, respectively. We refer to these settings as low and high density conditions or treatments throughout. Larval densities were chosen to vary per capita nutrition available in the two treatments. The chosen values are comparable with larval densities in natural habitats [51]. Previous studies involving similar larval densities identified direct and indirect effects of intraspecific larval competition on larval and adult traits of mosquitoes, including body size [52].

For all experiments, the stock larval nutrient medium was prepared at 3.3 mg/mL using standard fish food (Hikari Tropical First Bites, Petco, San Diego, CA, USA), incubated at 26°C for 24 hours, and used to prepare 12.5% stock dilution. Eggs were synchronously hatched in deionized (DI) water using a vacuum chamber and transferred to 300 mL of nutrient medium stock dilutions. Freshly hatched mosquito larvae in 12.5% dilution of the nutrient broth were housed in an incubator at 26±0.5°C and 70±10% relative humidity with a 14:10 hour day-night cycle. Mosquito larvae were monitored every 6 hours until pupation. Upon pupation, they were transferred to individually labeled vials containing water and monitored until emergence using a locomotor activity monitor (Trikinetics, LAM25). The activity monitor contains three infrared beams bisecting each vial just above the water level, repeatedly recognized by opposing infrared detectors. The LAM25 software recorded infrared beam breaks at repeated 60 second intervals to detect and record emerging mosquitoes. Modifying the locomotor activity monitor setup which otherwise is generally used for tracking mosquito activity allowed us to accurately quantify the development time of individual mosquitoes.

We recorded larval development times and proportion surviving through each molting event (four larval instars and pupa) and the metamorphosis into adults. The experiment on development time for stages consisted of five replicates in low density and five replicates in high density. Each day, all larvae were identified for stage and counted. On the day when an individual emerged as an adult, it was sexed using morphological features such as the structure of the antennae [53].

To determine the body mass, we conducted 13 replicates in low density and 8 replicates in high density. The mass of adult females, but not males, was recorded, resulting in measurements for a total of 120 and 158 adult females from low and high density conditions, respectively. Each adult female was weighed on emergence, and her total development time was recorded (note that in this experiment, development time of each larval stage was not recorded). Emergence time in body mass data was measured in hours; however, as our model uses a time step of one day, we divided emergence time by twenty-four and rounded up to the next day.

We determined survival using all of the above replicates from both stage and mass data sets plus two additional replicates at low density. The two additional replicates were conducted similarly to the experiments in which we studied body mass, except mass was not measured (i.e. only the total number of male and female that emerged was recorded). Thus, the data to determine survival values consisted of 20 replicates in the low density treatment and 13 replicates in the high density treatment, resulting in a total of 520 and 1014 first instar larvae initially, of which 412 and 656 survived to emergence in low and high density conditions, respectively.

#### 2.1.1 Key Features of Data

We aim for our model to reproduce key features observed in our experimental data. In particular, the model should differentiate timing of individuals based on sex and body size in different larval environments. In Fig 1a, we present data on development time aggregated by sex and density treatment. On average, males emerge slightly before females, and individuals in the high density environment exhibited greater variance in development time. Using a Welch *t*-test, we found a significant difference in development time for males and females (*df* = 192.92, *p*-value = 8.252e – 12 for low density and *df* = 300.53, *p*-value < 2.2e – 16 for high density). When we considered experiments by sex and density treatment (Fig 1c), we noticed that density impacts female survival: the average female survival in the high density treatment is approximately two-thirds of the low density treatment. Males showed a similar trend but to a lesser extent. With a *t*-test comparing between density treatments for each sex, we found that the difference in low and high density is statistically significant for females but not for males (*df* = 30.131, *p*-value = 0.001364 for females and *df* = 28.35, *p*-value = 0.1196 for males)

**Fig 1.**
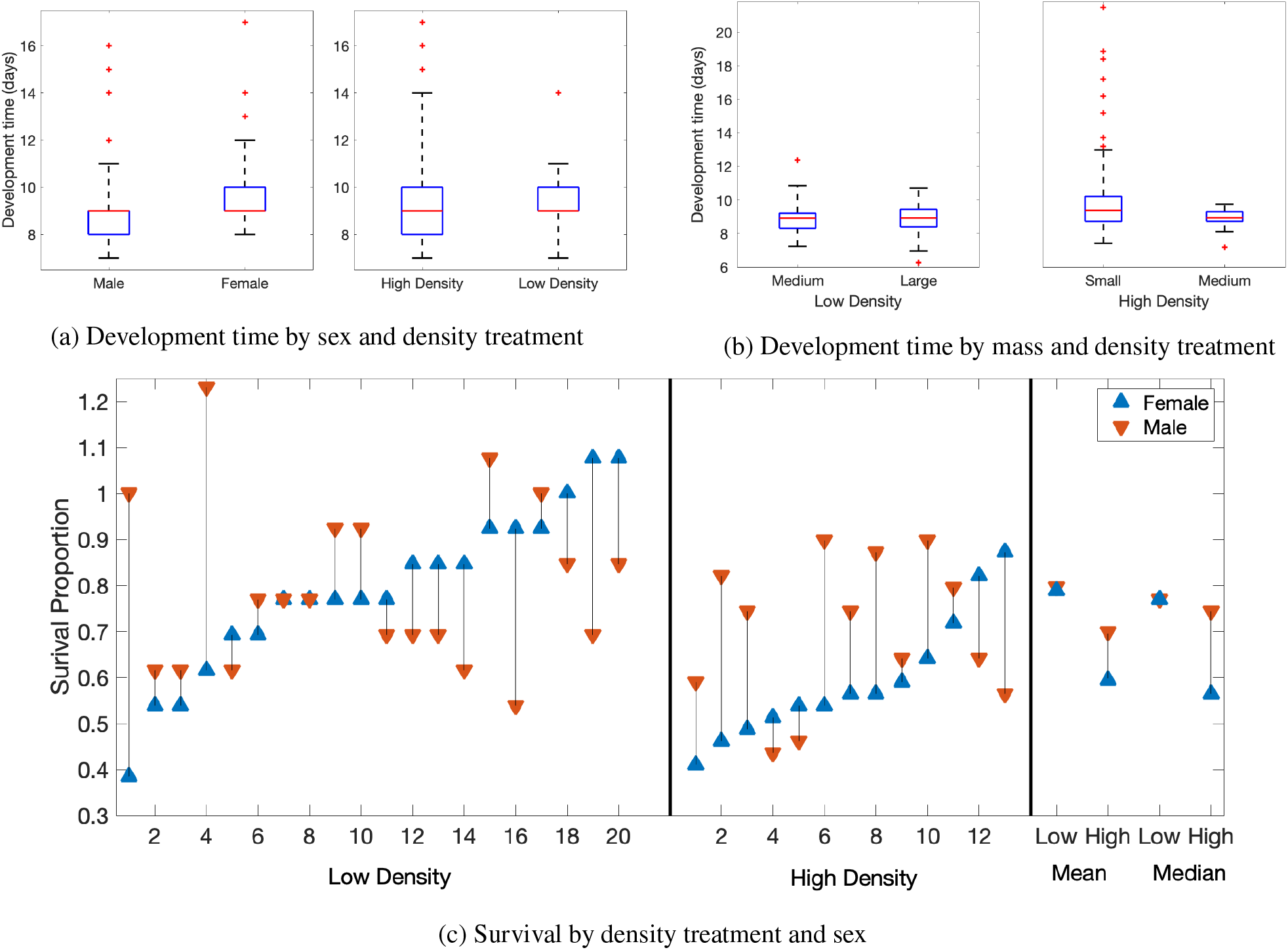
Experimental data on development time and survival. (a) Development time, in days, of mosquitoes across all experiments, categorized by (left) sex and (right) density treatment. (b) Development time by mass group and density treatment. Box plots of each mass group emerging under (left) low density and (right) high density treatments. This includes mass data, but excludes stage data. In (a)-(b) red lines represent the median of the data; the blue box indicates the upper and lower limits of the interquartile range (IQR); whiskers indicate 2*IQR; and red plus signs represent outliers. (c) The proportion of mosquitoes which survive by density treatment and sex for each replicate of the low (left) and high (middle) density treatments. (right) The mean and median survival proportion for all experiments by density. In (c), blue triangles represent females and red triangles males. The experiments are sorted by female survival proportion, where we assume 50% males and 50% females initially. As individual replicates may differ in their initial sex percentages, some calculated survival proportions are greater than one.

In our analyses, we considered three general mass groups: small (less than 1.5 mg), medium (between 1.5 and 2.5 mg), and large (greater than 2.5 mg). These three groups were chosen by dividing the data from mass at emergence, which ranged from 0.69 to 3.35 mg, into approximately equal thirds. There is a noticeable difference in the proportion of each mass group emerging based on the environmental conditions (Table 1). In both density conditions, food was only provided at the beginning of the experiment. In the low density treatment, this food was sufficient for all larvae over the time span required for development. Thus, the majority of mosquitoes that emerged are of the large mass group and only a single mosquito emerged in the small mass group. In the high density treatment, the amount of food was likely insufficient for successful development of all mosquitoes. As a result, none of the mosquitoes emerged in the large mass group and the majority were in the small mass group.

**Table 1.** Mosquito emergence by body mass in low and high density conditions. The number (percentage) of mosquitoes in each body mass group: <1.5 mg (small), 1.5-2 mg (medium), and >2.5 mg (large) divided by environmental conditions: low density and high density.

| Environment | Small | Medium | Large |
| --- | --- | --- | --- |
| Low | 1 (<1%) | 24 (20%) | 95 (80%) |
| High | 114 (77%) | 44 (23%) | 0 (0%) |

In Fig 1b, we show data on development time by mass group for each of the density treatments. There is a significant difference in the development time of the small mass group compared to the medium group in the low density treatment (*p* = 0.012, *n* = 158). No differences between mass group are seen in the high density treatment. The data does not meet the normality assumptions necessary for ANOVA; thus, we performed a Kruskal-Wallis and Dunn test to determine if, and which groups, are different. Although the small mass group is significantly different, the Kruskal-Wallis test effect size is small (0.046), so we do not consider a variation based on this difference.

### 2.2 Model

We developed a discrete time model of an *Aedes* mosquito population which incorporates each biological stage of the mosquito’s life cycle: four larval instars, pupa, and adult. In our model, we include effects of larval density and resources on development time and growth (as measured by mass). To that end, we divided each of the larval and pupal stages further by mass, which we denote using a subscript: small (*s*), medium (*m*), and large (*l*). Our determination of the mass range for each of these categories is as described in section 2.1.1.

The time step for our model is one day. We assume all individuals begin in the first instar larvae (*L*1) as our data does not include eggs. Individuals move through successive larval stages (*LX*, where *X* denotes the instar: 1, 2, 3, or 4). The proportion of juvenile mosquitoes that develops each day is governed by the function *f*(*N*), where *N* is the total number of larvae at time t. We assume that development of juvenile mosquitoes is density-dependent [54,55], and let the density in the model equal the current larval population (*N*). See details on *f*(*N*) in section 2.2.2. At each transition, a proportion of individuals die, which we denote as *μ*(*N*). See details on *μ*(*N*) in section 2.2.3. We allow individuals to move first and then introduce mortality.

All mosquitoes begin in the *L*1 stage to mimic the initial condition of our experiments. We assume that, in each replicate, 50% of the eggs are male, and 50% are female. *β*(*t*) represents the number of L1 larvae of a given sex inputted in the system at time t. For all data fitting, *β*(*t*) = 0 when *t* ≠ 0. For model simulations of low density treatment, *β*(0) = 26/2, and of high density treatment, *β*(0) = 78/2. During the larval stages, mosquitoes may remain in their current mass group or move to a higher mass group. We assume that mosquitoes changing mass groups can only move to the next highest mass group in a time step (i.e., a small mosquito cannot become a large mosquito in a single day). For example, during the transition from the first larval stage (*L*1) to the second larval (*L*2) stage, individuals can either stay in the small mass group or move to the medium mass group. Similarly, each individual in L2 that moves to the third larval stage (*L*3) can stay in the same mass group or move to a higher mass group. We note that once mosquitoes enter the large mass group, they cannot grow larger in mass. During the pupal stage individuals do not eat, so they remain in their current mass group. A diagram depicting our model is shown in Fig 2a. At time step *t*, the growth functions, *G_i_*, determining movement to a higher mass group depend on resource (*r*) and larval density (*N*). The proportion of individuals that grow from small to medium mass is given by *G*_1_(*r*/*N*), and *G*_2_(*r*/*N*) determines the proportion of individuals that grow from medium to large mass. See details on *G_i_*(*r*/*N*) in section 2.2.4.

**Fig 2.**
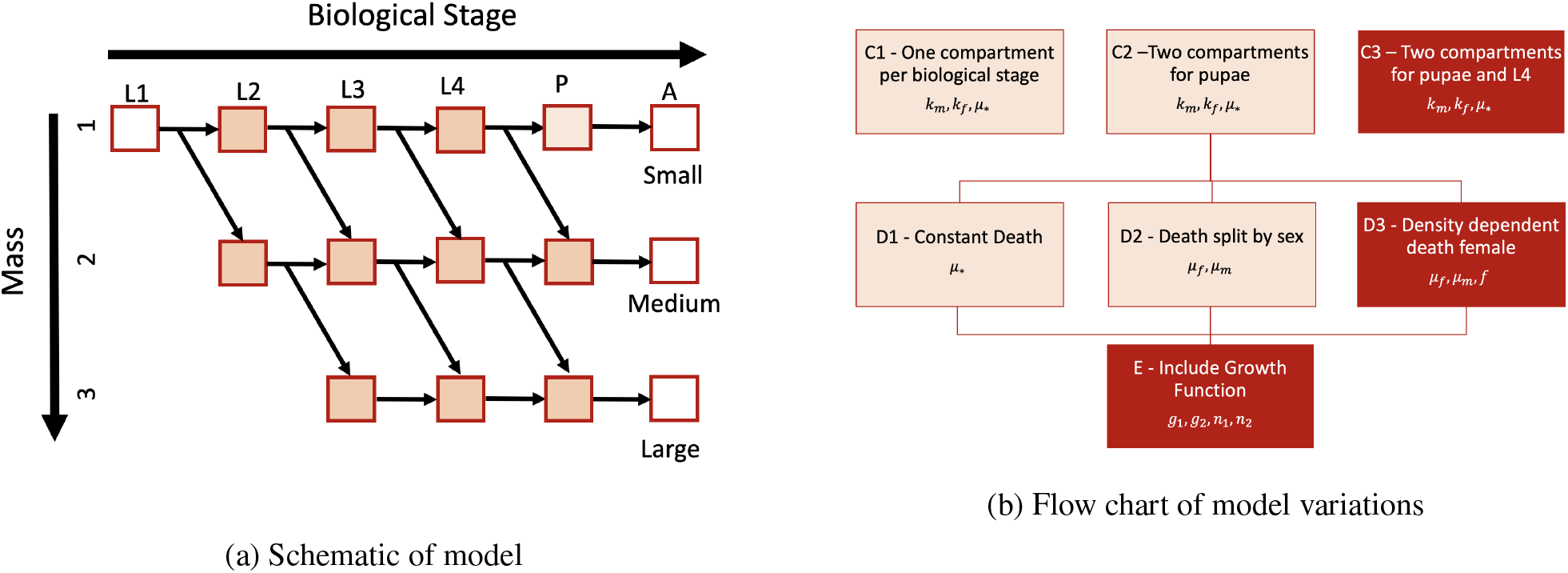
Model formulation and variations. (a) Schematic of our mathematical model describing stage and mass. As an individual moves horizontally (age axis), it advances to a later biological stage from larval stages 1 to 4 to pupae to adult. As an individual moves vertically (mass axis), it grows larger in mass. (b) Flow chart of each variation considered. The first variation (top row) is in regards to the number of compartments. We include 5, 6, or 7 total compartments, in C1, C2, and C3 respectively. The second variation is in mortality: single constant death (D1), two constant deaths for each of male and female (D2), and a density dependent death for females and constant death for males (D3). The third variation is the inclusion of a growth function, E. The darker boxes are the version of each variation chosen as best based on the AIC. See section 2.4 for description of the fitting.

The system of equations describing our model is given below. We do not track adult populations, but only record the emergence of adults by sex and mass group through time. For example, *F_s_*(*t*) represents the number of newly emerged females in the small mass group at time *t*, and *M_l_*(*t*) represents the number of newly emerged males in the large mass group at time *t*. Note that for the larval and pupal stages, we track males and females separately to account for differences in development and survival that are sex-dependent, but we only present larval and pupal equations for a single sex here as the equations are structurally identical. While the equation structure is identical, the following functions differ depending on sex: death proportion (*μ*(*N*)), the proportion which develop (*f*(*N*)), and the proportion which grow (*G*_1_(*r*/*N*),*G*_2_(*r*/*N*)), where *N* is total larvae (males + females) and *r* is resources.

Equations for the aquatic phases (identical for both male and female) are given by:

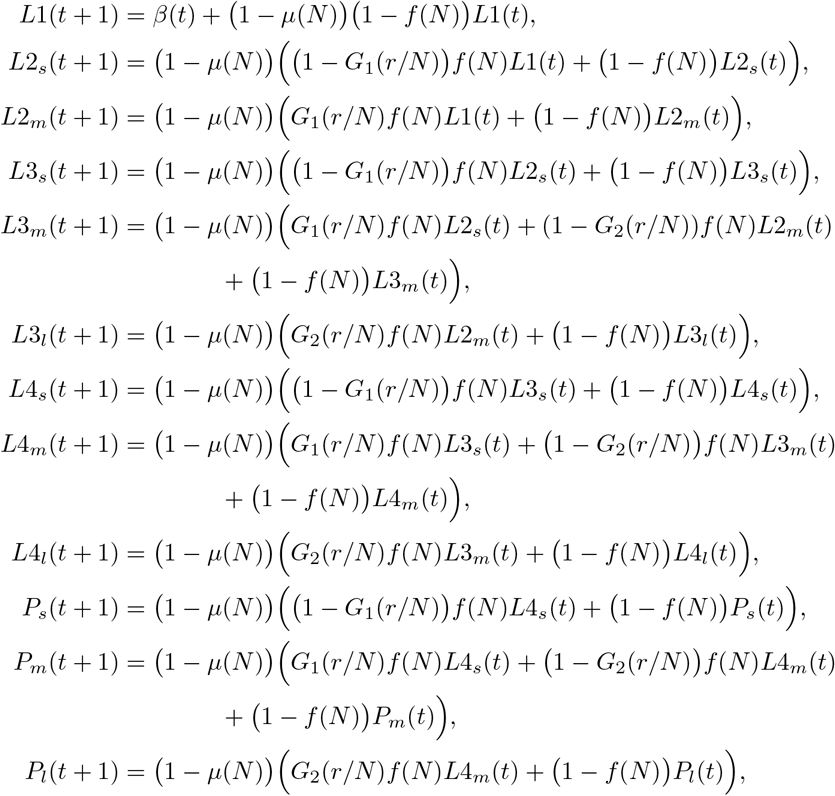

and for emerging male adults:

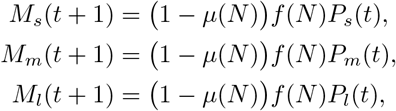

and for emerging female adults:

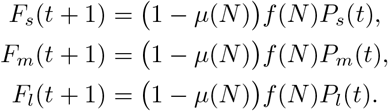

#### 2.2.1 Resources

We incorporate the amount of available resources (i.e. food) to larvae in our functions for growth and development. We assume that there is a fixed maximum amount of food necessary for each larvae and any excess food does not help or hinder larval growth or development. The amount of food (*r*) that is sufficient for a single larva occurs when *r* = 1. If there are *N* larvae, *r* = *N* is the necessary amount of food for all larvae.

However, we assume that when there are lower levels of food, for example, as the result of decreasing resources, larvae experience slower growth. Thus, we consider growth a function of food per larvae, 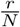, where *r* is the resource and *N* is the current total larvae.

We assume that resources decay such that each day,

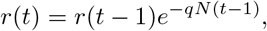

where *N*(*t*) is the total larvae at time *t* and *q* is the rate of decay. The value for *q* is chosen such that the resources available in the low density treatment would be sufficient for all larvae while the resources available in the high density treatment would run out before all larvae pupate. The value for *q* was fixed to 0.01. In section 3.4.1, we discuss our choice for *q* in more detail and include a sensitivity analysis.

#### 2.2.2 Development time

In previous work, we showed that development time is affected by density [55], which was also noted by others [54]. To model density-dependent development time, we incorporate the function *f*(*N*), which is the proportion of individuals that develop to the next life stage. For *f*(*N*), we use the Maynard-Smith-Slatkin density function, which is the best fit formulation from [55], and is given by

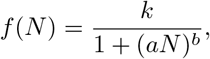

where *k* is the maximum development rate, *N* is the total current larvae, and *a* and *b* scale the importance of density. We use the parameter choice *a* = 0.0043 and *b* = 1.61 found in previous work [55]. Here, we employ different *f*(*N*) for males and females and fit two different values for the maximum development proportion, *k*: one for males, denoted *k_m_,* and the other for females, denoted *k_f_*.

#### 2.2.3 Death proportion

A proportion of individuals from each compartment die each day. In an initial pass, the death proportion for each day is taken directly from the experimental data. In this case, we compare each replicate separately and use the proportion of individuals that die each day from that specific replicate. We use the daily death proportions by replicate to fit the development time. After we fit the daily maximum development proportion (*k_m_, k_f_*, see section 2.4.1), we combine data on mortality from all replicates and fit a single constant for death proportion.

We then consider a density-dependent death function for females. In the experiments, we observed that females had greater survival in low density compared to high density (Fig 1c). While males showed a similar trend as females, the difference observed between low and high density treatments was not statistically significant. Thus, we do not consider density dependent mortality for males. To incorporate density dependence into our death function for females, we use a Hill function, given by

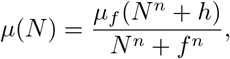

where *n* = 3 is the Hill exponent, *h* and *f* are constants such that *h* < *f* and 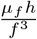 is the minimum proportion of individuals that die and *μ_f_* is the maximum proportion. The inclusion of the *h* in the numerator, a departure from a traditional Hill function, allows the lower bound of the function to be greater than zero. If *h* equals zero, this returns the traditional Hill function. We fix *h* = 100, so that the lower bound is 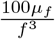. See section 3.4.4 for more discussion and a sensitivity analysis of *f*.

#### 2.2.4 Growth

The growth functions determine the proportion of mosquitoes that move from one mass group to the next largest mass group. For implementations of the model that do not involve mass (sections 2.4.1-2.4.2), we assume the growth functions are zero. In such cases, nothing depends on mass so there is no need to track which the mass group of mosquitoes in a particular life stage. In all other cases, we use our fit to the functional form below for *G*_1_ and *G*_2_. At each time step, the function *G*_1_ determines the proportion of individuals that grow from small to medium mass

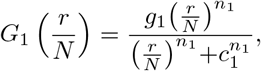

and *G*_2_ those that grow from medium to large mass

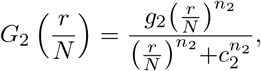

where *n*_1_ and *n*_2_ are Hill coefficients; *c*_1_ and *c*_2_ are the value where half maximal growth occurs; and *g*_1_ and *g*_2_ are the maximum growth proportions. Note that *G*_1_ and *G*_2_ inherently depend on time as *r* and *N* change with time.

The proportion which grow from one mass group into a higher mass group is reduced when 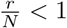. Furthermore, in the absence of food (*r* = 0), the proportion which grows is defined to be zero. If the amount of food is well above that necessary for the current number of larvae, i.e. 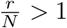, the proportion which grows does not exceed *g_i_* ≤ 1. As *r* = 1 is sufficient food for one larva, we expect the growth function to reach the maximum approximately when 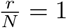. Thus, we assume that half maximal growth occurs when 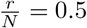, i.e. *c*_1_ = *c*_2_ = 0.5. See section 3.4.3 for more discussion and a sensitivity analysis on *c*_1_ and *c*_2_.

The parameters *g*_1_, *g*_2_, *n*_1_ and *n*_2_ are determined to be consistent with the resulting mass difference observed under low and high density treatments. We assume that growth to the large mass group is more harshly affected by lack of resources than growth to the medium mass group. Furthermore, we consider different growth functions by sex, such that maximal growth proportion differs for males and females. Specifically, we assume that a lower proportion of males grow, and let maximum male growth be *νg*_1_ and *νg*_2_ where 0 < *ν* ≤ 1.

### 2.3 Model Variations

In our base model, we assume a single constant daily death proportion for both males and females (*μ*), but have different daily maximum development proportions for males (*k_m_*) and females (*k_f_*). As we are not initially incorporating data on mass, we set maximum growth proportions for both males and females to zero, *g_i_* = 0.

In our analysis, we consider three sets of variations to the base model: variations in the number of compartments (denoted by C); variations in death proportions (denoted by D); and inclusion of the growth function (denoted by E). A diagram of the variations is shown in Fig 2b.

#### 2.3.1 Compartment Variations: C1-C3

First, we altered the total number of compartments, which resulted in three different variations of the model. To begin, we allow for each aquatic stage to encompass only a single compartment in the version denoted C1. We then add a second compartment to the pupae stage, so that there are two compartments for each sex and body size of pupae in the version denoted C2. Finally, we split apart both pupae and L4 into two compartments, instead of a single compartment for each, in the version denoted C3. The addition of these compartments forces the minimum development time to be longer for all mosquitoes. The choice of including the compartments in L4 and pupae is motivated by observations on the time to emergence in our experiments (Fig 1c). It is important to note that the additional compartments only extend the development time but do not impact growth as no growth can happen during transition between the first and second sub-compartments within L4 or during the pupal stages. In our model, we denote the second compartment in each of L4 and pupal stages with an asterisk. For example, *P_l_* is the first compartment for pupae in the large mass group and *P_l_*\* is the second compartment.

Finally, as detailed in section 2.4.1, we compare the results obtained from the three variations and choose the variation that best fits larval timing and adult emergence. Once we determine the optimal number of compartments for the model structure, we use this as the starting point to consider variations in how we model death.

#### 2.3.2 Death Variations: D1-D3

Next, we consider different ways to incorporate death by studying three variations with different assumptions on mortality. The death proportion is initially a single constant value for both males and females in the version denoted D1. The second version, denoted D2, uses a different constant proportion for each sex. Finally, we consider a density-dependent death function for females, but not males, in the version denoted D3. We do not include a density-dependent death function for males as the data did not support differences for the males by density treatment. We compare all three versions of the incorporation of mortality as described in section 2.4.2. We use the best fitting model to consider inclusion of the growth functions.

#### 2.3.3 Growth Function Included: E

After determining the number of compartments and the form for the death function, we fit the two growth functions, *G*_1_ and *G*_2_. At each time step, these functions determine what proportion grows from small to medium mass and from medium to large mass, respectively. Details of the growth functions are found in section 2.2.4. We denote this as variation E.

### 2.4 Fitting Parameters

We describe our fitting for each variation of the model. Throughout, we use 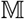 to refer to model output and 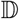 for experimental data. We list all parameters with their description, the standard value, and variations considered in Table 2.

**Table 2.** Model parameters. For each parameter, we include its representative symbol, a description, the standard value used, and the range of values considered. For parameters marked fixed, we used a single value during the fitting process, but performed univariate sensitivity. For parameters marked fitted, the range listed under variation is the constraint when fitting using MatLab function `fmincon`.

| Symbol | Description | Choice | Value | Variation |
| --- | --- | --- | --- | --- |
| $q$ | Decay rate of resources | Fixed | 0.01 | 0.0005-0.05 |
| $a$ | Exponent on density in development time | Fixed | 0.0043 | 0.001, 0.01 |
| $b$ | Coefficient on density in development time | Fixed | 1.61 | 1, 2 |
| $k_m$ | Maximum male daily development proportion | Fitted | 0.94 | 0-1 |
| $k_f$ | Maximum female daily development proportion | Fitted | 0.85 | 0-1 |
| $\mu_m$ | Maximum male daily death proportion | Fitted | 0.025 | 0-0.5 |
| $\mu_f$ | Maximum female daily death proportion | Fitted | 0.066 | 0-0.5 |
| $f$ | Female density dependent death parameter | Fitted | 24 | 1-100 |
| $h$ | Constant in density dependent death | Fixed | 100 | 1-2000 |
| $n$ | Exponent in female density dependent death | Fixed | 3 | 1-10 |
| $g_1$ | Maximum daily growth proportion from small to medium | Fitted | 1 | 0-1 |
| $g_2$ | Maximum daily growth proportion from medium to large | Fitted | 0.46 | 0-1 |
| $n_1$ | Exponent of growth function from small to medium | Fitted | 3 | 0.5-12 |
| $n_2$ | Exponent of growth function from medium to large | Fitted | 9 | 0.5-12 |
| $c_1$ | Per capita resource for half maximal growth for small to medium | Fixed | 0.5 | 0.3, 0.5, 0.69 |
| $c_2$ | Per capita resource for half maximal growth for medium to large | Fixed | 0.5 | 0.3, 0.5, 0.69 |
| $\nu$ | Male proportional growth | Fixed | 0.75 | 0.5-1 |

#### 2.4.1 Estimating Development Time: Variations C1-C3

In our compartment variations, we ultimately fit three parameters (*k_m_, k_f_, μ*(*N*) = *μ*_*_) using a two step process. We began by simultaneously fitting the maximum development proportion for males (*k_m_*) and females (*k_f_*). In order to separate effects of density on death and on development time, we calculated the daily death proportion directly from each replicate by dividing the number of individuals that died on the previous day by the total number of larvae and pupae present on the previous day.

We fit the maximum development proportion for males and females by minimizing a summed squared error of the difference in total larvae time and the total emergence of adults from each data set. There are ten sets of data: five replicates in high density and five replicates in low density.

Let the 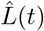 be the vector of the sum of each aquatic stage (across mass groups) at each time *t* given by

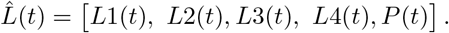

Let the total sum of the aquatic stages from a given model simulation be 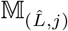, where *j* ∈ {*L, H*} represents low and high model density treatments. The first subscript of 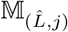 indicates element-wise sum over 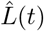.

Let the total number of emerged adult females from a given model simulation be 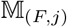 for the *j*th data set. The total number of emerged male mosquitoes are represented similarly by 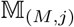. Thus, model output for emerged adults is given by

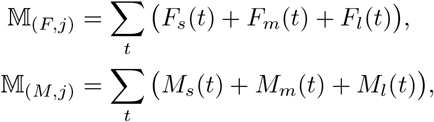

where *M_i_*(*t*) and *F_i_*(*t*) for *i* ∈ {*s, m, l*} are the number of emerging males and females, respectively, of a given mass group at time *t*. While 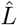 is a vector, 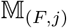 and 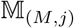 are each scalars. The data from a particular replicate *j* is represented similarly but with 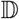 rather than 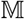.

We fit the development proportions by finding the square of the difference between model output and data, weighting each term, and then summing across all replicates. Specifically, our error formula is given by

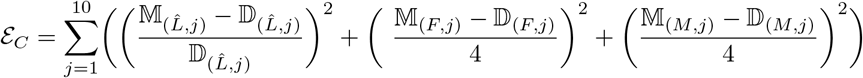

where the division and the square occur element-wise in the first term. We chose the weights in the error formula so that the data on the emergence of adults (second and third term) has more weight than data on time spent in the biological stages (first term). We placed more weight on the adult results as sex is not separable in the data until the adult stage.

To find the minimum error *ε_C_*, we used `fmincon` in MatLab allowing both *k_m_* and *k_f_* to be constrained between 0 and 1. The MatLab function `fmincon` uses an interior point algorithm which attempts to solves the Karush–Kuhn–Tucker (KKT) conditions for an approximated problem. If this step is not possible, then it uses a trust region conjugate gradient step. We choose a nine by nine grid (values between 0.1 and 0.9 incremented by 0.1 for each of *k_m_* and *k_f_*) of initial starting points.

Once we determined optimal *k_m_* and *k_f_* in our first step, we used these values in estimating a single daily death proportion, *μ*(*N*) = *μ*_*_, for all replicates. In this case, the only difference between model output from high (*H*) and low (*L*) density treatments is the initial number of mosquitoes in L1, our initial condition. We found a constant *μ*(*N*) = *μ*_*_ that minimizes the death error, *ε_D_*, found in section 2.4.2. We used appropriate weighting for low and high density with 5 replicates for each condition.

Overall, we fit three parameters: *k_m_, k_f_*, and *μ*_*_. In the first step, we fit *k_m_* and *k_f_* simultaneously. Then in the second step, we used these values when we estimate *μ*_*_.

#### 2.4.2 Estimating daily death proportions: Variation D1-D3

For our death variations (D1-D3), we fitted parameters related to death proportion (*μ*_*_, *μ_f_, μ_m_, f*) and calculated the total number of males and females that emerged. The data consists of 13 replicates in high density and 20 replicates in low density. For these replicates, we used the total number of males and females that emerge as adults.

Let the total number of emerged adult females from the model in low density be 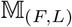 and in high density as 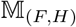. Similarly, the total number of emerged adult males in low and high density is 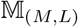 and 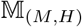, respectively. Thus, our model output is given by

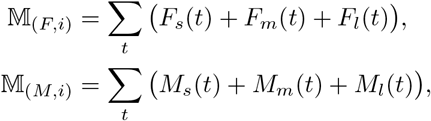

where *i* ∈ {*L, H*}. Similarly, let the total number of females from the kth replicate in low density be given as 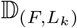 and from jth replicate in high density as 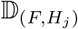.

We determined the sum of the squared difference between data and model output of the total number of emerged males and total number of emerged females in each density treatment, as given by

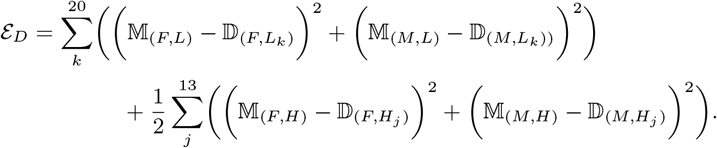

Note that we scaled the sum for the high density treatments by one-half as we aimed for approximately equal weighting for data from both high and low density treatments. Recall there are a total of 20 replicates in low density, each assumed to start with 13 individuals of each sex, and 13 replicates in high density, each assumed to start with 39 individuals of each sex. Thus, there are approximately twice as many total larvae across all high density replicates, so we scale the sum by one-half.

In order to minimize our error *ε_D_*, we used MatLab function `fmincon` on a range of initial values for the parameters. The precise parameters that we fitted was dependent on the death proportion variation considered. In variation D1, there we fitted a single death constant (*μ*(*N*) = *μ*_*_). In D2, we fitted two death constants, one for males (*μ*(*N*) = *μ_m_*) and one for females (*μ*(*N*) = *μ_f_*). Finally, in D3, we fitted two non-fixed parameters (*μ_f_, f*) for a density dependent death function for the females 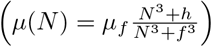 and a single constant death (*μ*(*N*) = *μ_m_*) for the males. See section 2.2.3 for more details on the density dependent death function. In our fitting, the values *μ_f_, μ_m_*, and *μ*_*_ are constrained between 0.001 and 0.5, and *f* is constrained between 1 and 100.

#### 2.4.3 Estimating Growth Function Parameters: Variation E

In variation E, we investigated mass-dependent growth. As previously noted, we split individuals by body mass into three groups: less then 1.5 mg (small), between 1.5 and 2.5 mg (medium), and greater than 2.5 mg (large). To determine the growth function, we fit the parameters *n*_1_, *g*_1_, *n*_2_, and *g*_2_ as described in section 2.2.4 and fix the resource decay rate to *q* = 0.01 as described in section 2.2.1. The latter results in complete resource usages in the high density treatment prior to the emergence of all individuals as adults, but allows resources to remain in the low density treatment even after all individuals emerged.

We have less data on mass size for males. Thus, we first found the average development time for females and males separately. Then, we determined *ν* to be the relative proportion that males grow compared to females, by dividing the male average with the female average. See section 3.4.2 for more discussion on *ν* and a sensitivity analysis.

The data is comprised of 13 replicates in low density and 8 replicates in high density. For each replicate, we use the total number of emerged adult females in each mass group. Recall that *F_s_*(*t*) (*F_m_*(*t*), *F_l_*(*t*)) are the small (medium, large) females that emerge as adults at time *t*. Let the total number of small females that emerged in the *k*th replicate be given as 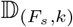 (similarly, 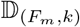 and 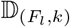 for medium and large, respectively). The model output for the total number of small females emerged is given by 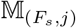 (similarly 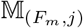 and 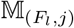 for medium and large, respectively) where *j* = *L* for low density and *j* = *H* for high density. In order to fit the parameters for the growth functions, *G*_1_(*r*/*N*) and *G*_2_(*r*/*N*), we minimized the squared difference between the proportion of each mass group (small, medium, and large) in the data compared to the model run. The error function is given by

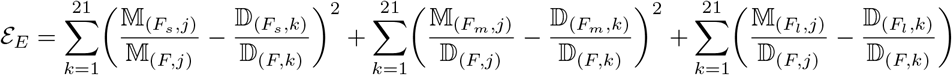

Note that the value for *j* in the subscript of 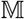 is determined by the specific data replicate considered: if it is low density, then *j* = *L*, and if it is high density, then *j* = *H*.

In order to find the minimum error *ε_E_*, we used the function `fmincon` in Matlab with over 150 initial value choices. We constrain *g*_1_ and *g*_2_ between 0 and 1, and *n*_1_ and *n*_2_ between 0.5 and 12.

### 2.5 Model Comparison

We have three different sets of variations of the model we fit: C1-C3, D1-D3, E (Fig 2b). We compared each version within its variation group (row in Fig 2b). We use the Akaike information criterion (AIC), a common metric to compare between models, to choose the best-fitting model. The AIC is a relative measure of the maximum likelihood of a model that deducts for the complexity of the model by reducing by the number of parameters needed. In particular, we use a second order biased correction, AIC_*c*_, as the sample sizes in our data are relatively small. We assume errors are normally distributed and use different versions of least squares error from the data (*ϵ_i_*) described above. The AIC_*c*_ we employ is given by

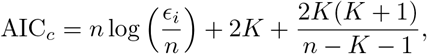

where *K* is the number of parameters in the model, *ϵ_i_* is the error as described above, and *n* is the sample size of the data [56].

For the first variations (C1-C3), we varied the number of compartments, but the number of parameters being fitted does not change. Since there is no complexity difference in the number of parameters, the AIC_*c*_ simply compares the errors without any difference due to parameters. After determining the maximum male and female development proportions, these parameters are fixed for the next set of variations.

Next, we determined the best representation of death, D1-D3. The number of parameters in these versions varies from one in the simplest case to three in the most complex case. Using the AIC_*c*_ we determined which version is best and used that version when we estimated parameters in the growth function.

Finally, in variation E, we fit parameters in the growth functions by estimating four parameters. We only considered a single mathematical formulation of the growth functions as mass-dependent growth of larvae is a poorly understood process. Furthermore, our growth functions do not mechanistically describe growth but only represent the proportion that emerge as a larger mass.

## 3 Results

### 3.1 Development Time Fit: Variation C1-C3

We observed that the predicted adult emergence in the model generally occurred at similar times to that in the data. The model, however, spreads emergence across multiple days compared to the data, which tends to sharply peak across one or two days. This results in the model output showing lower peaks on any given day. In the absence of extra compartments in L4 and pupae, the model results show the timing of emergence of both males and females in either density treatment begins two to three days earlier in the model than observed in the data (Fig 3a, dashed dotted light blue line). Adding in a second pupal compartment reduced this difference to one to two days (Fig 3a, dashed maroon line). Finally, adding in a second compartment to the L4 stage in addition to the second compartment for the pupal stage further reduced this difference (Fig 3a, solid blue line). Emergence in high density started on the same day in the model output and in the data, and emergence in the model output under low density started one day earlier. In all versions of the model, the model predicted emergence spread across a wider range of days. In particular, the model suggests a longer tail of later emergence times than in the data Fig 3a).

**Fig 3.**
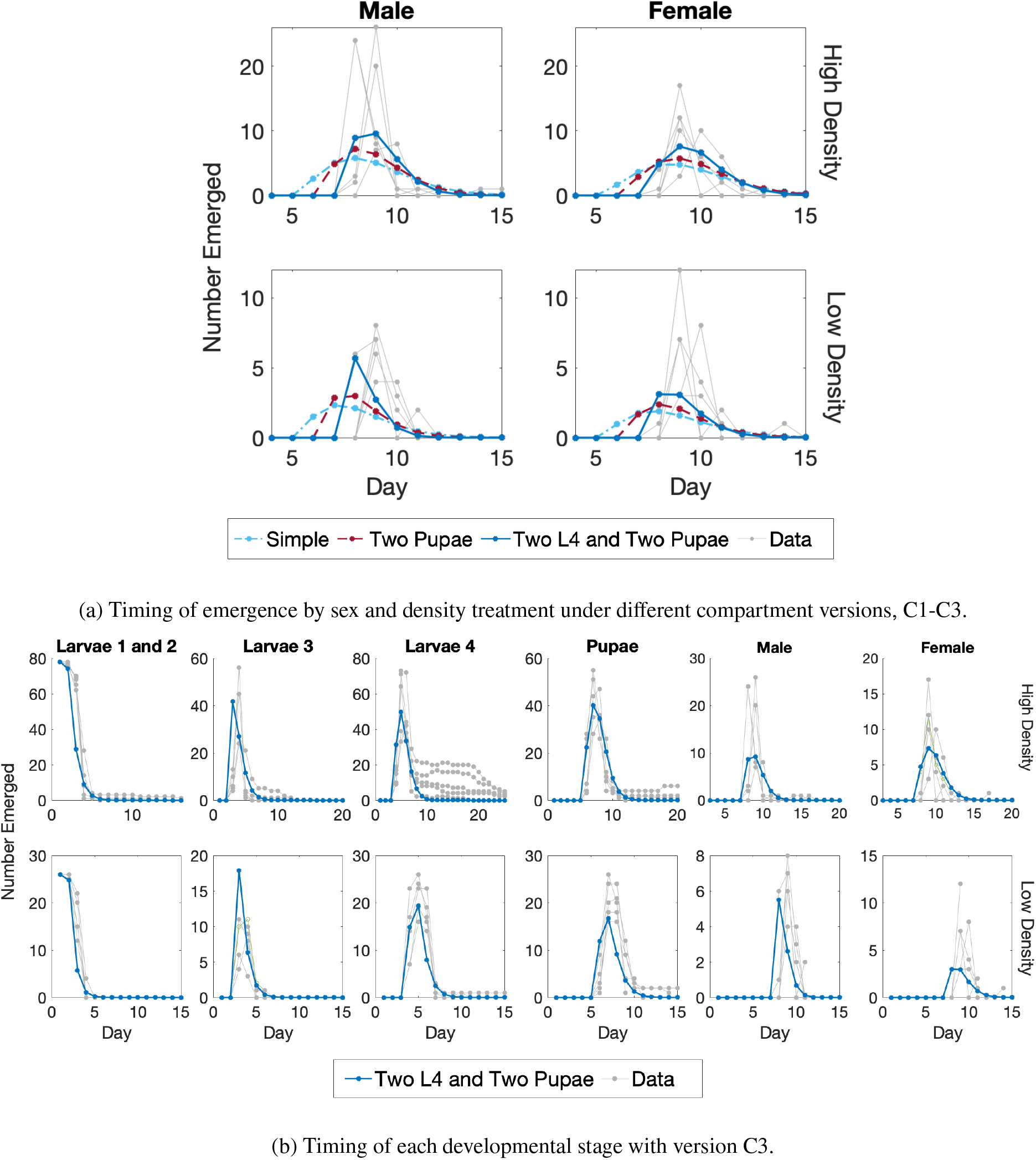
Fitting results for compartment versions, C1-C3. (a) Data and model output in the high density (top row) and low density (bottom row) treatments. The results are split by sex: males (left column) and females (right column). Each panel shows the number of adults that emerged on a given day. The gray lines each represent an individual experimental replicate. For the model, the only difference between low and high density treatments is that the initial values are different. The best fit for: C1, the base model with only one compartment per biological stage (dashed dotted light blue line); C2, including a second compartment for pupae (dashed maroon line); and C3, including second compartments for both pupae and L4 (solid blue line). (b) Model output in version C3, where both L4 and pupae have two compartments, in high density (top row) or low density (bottom row) treatments. From left to right, the panels show: combined L1 and L2 stages; L3 stage; L4 stage; pupae; emerging males; and emerging females. Each grey line is an experimental replicate.

As we employ a constant death rate (rather than time varying) in the model, we expect slightly more males than females as they develop faster. Even with a shorter development time for males, we found that the model predicts that the number of males and females emerging are nearly identical when assuming identical death rates. Experimentally, in the high density treatment 17 - 35 (mean 28.2) males and 16 - 25 (mean 20.2) females emerged. In contrast, in the model version approximately 26 males and 25 females emerged. In the low density treatment, 8 - 13 (mean 10) males and 7 - 13 (mean 10.4) females emerged in the experiments. While in the model, approximately 9 females and 9 males emerged. The model results are approximate as fractions of individuals can be represented so we round to the closest whole number.

Using the AIC_*c*_ value to compare the model output with the data on the development time, we found that all versions of the model are remarkably similar. We found AIC_*c*_ values of 22.9, 22.3, and 21.3 for the base model (C1), including a second compartment for pupae (C2), and including a second compartment for both L4 and pupae (C3), respectively. We found the model with the two extra compartments to be best among the three as it had the lowest AIC value. When we examined our results by individual stage separately, rather than by total development time, we saw that the model matches the data well at each stage (Fig 3b). It is clear, however, that the model does not capture all the features of each stage for individual data replicates, such as the persistence of individuals in L4 in high density. This is unsurprising as the data is more varied in the later stages, and the model fits the average of the data, not each replicate individually.

### 3.2 Death Proportion Fit: Variation D1-D3

We found that the model, with the best fit for each variation, reproduced the median value of the data well (Fig 4). For the death proportion versions, we obtained AIC_*c*_ of 59.6, 61.2, and 57.4 for D1, D2, and D3, respectively. Thus, version D3, with density-dependent death for females, is the best fit model.

**Fig 4.**
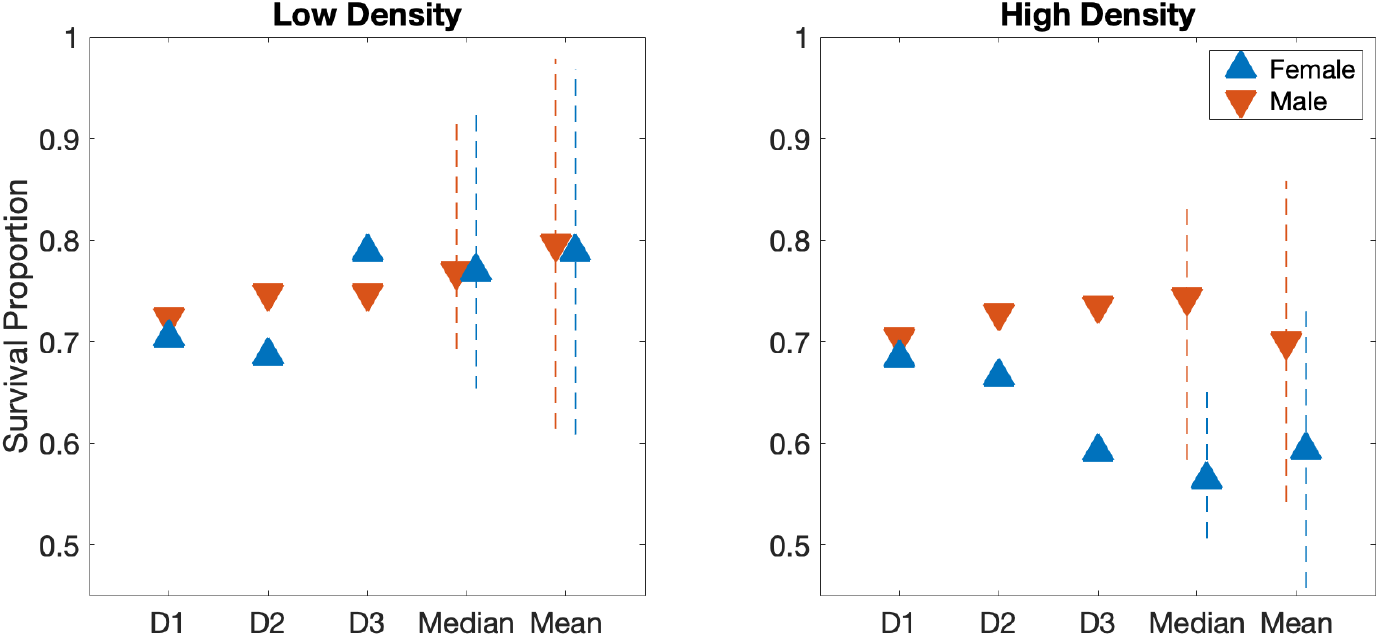
Total Survival proportion by sex and density treatment for experimental data and model versions D1-D3. Survival proportion of males (red) and females (blue). The model (D1-D3) under low density (left) and high density (right) treatments. On the far right of each panel, the values for median and mean of all data sets comprising of 13 high and 20 low density replicates, with dashed lines indicating one quartile above and below the median and one standard deviation around the mean.

In version D1, we fit a single constant for both male and female mortality across high and low density treatments. While we fit daily mortality, we discuss results in terms of overall survival, where survival can be determined approximately from mortality as: total survival proportion = (1 - mortality proportion)^development time^. Experimentally, we observed that while females in the low density treatment and males in both density treatments had a survival proportion near 0.75, the females in the high density treatment fared much worse. As all four situations are indistinguishable in D1, the single constant fit resulted in a survival proportion of 0.70. This is lower than the observed survival of 0.75 in three of the cases as it accounts for the lower survival exhibited by females in the high density treatment.

In D2, two parameters are included for mortality: one for males and one for females. Separating by sexes did not produce a better fit considering the added complexity of an extra parameter. In contrast, the inclusion of density-dependent death for females, as in version D3, allowed for different mortality by density condition for females. With this model version, we found a mortality proportion close to the median of the experimental data, and this version had the lowest AIC_*c*_ value. The total number of males that emerged in version D3 of the model is approximately 10 and 29 in low and high density, respectively, and the median of the number of total males in the data is 10 and 29 (low and high density, respectively). For females, the model outcomes for D3 were approximately 10 and 23 total females, and the median of the data was 10 and 22 (low and high density, respectively).

### 3.3 Growth Function Fit: Variation E

In the experiments, several low density replicates were almost entirely composed of large individuals (Fig 5a, right). In fact, nearly half of emerging adults in the low density replicates were entirely from the large mass group. In contrast, there were several replicates in high density treatment with only small individuals (Fig 5a, left). When we estimated parameters in our growth functions, the model fit to the mean of the experimental data, which is skewed by replicates with all individuals in a single mass group. This is particularly true for the large mass group in the low density treatment and the small mass group in the high density treatment.

**Fig 5.**
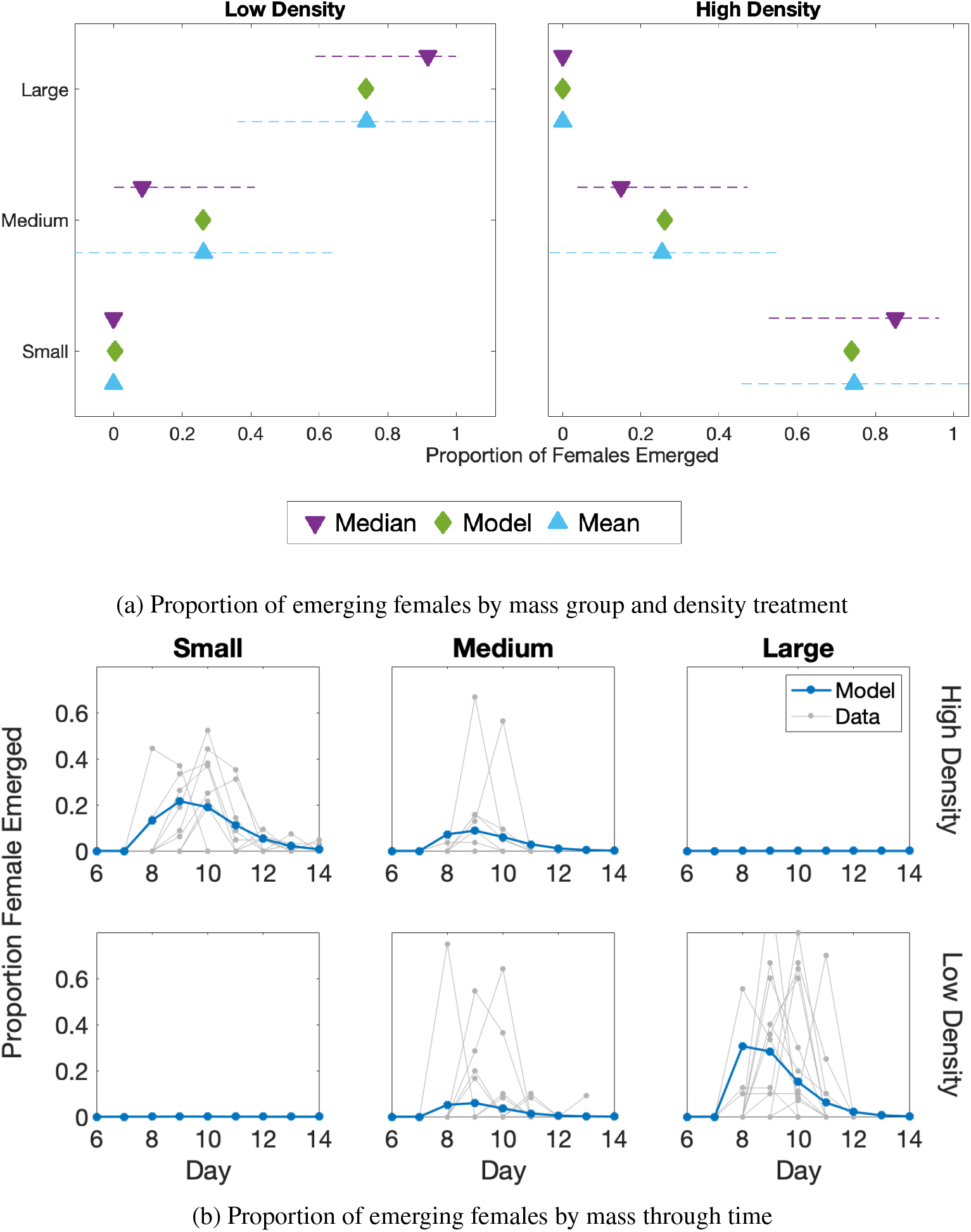
Fitting results for inclusion of the growth function, variation E. (a) The proportion of emerged females in a particular mass group is represented on the x-axis, and the mass group of these females on the y-axis. The green diamond is the best fit model. The median of the data is represented with a purple triangle with a dashed line for one quartile above and below. The blue triangle represents the mean of the data, and the dashed line is one standard deviation around the mean. The left panel is low density and right is high density. (b) Proportion of females that emerge from each mass group through time: small (left), medium (middle), large (right). The top row is high density and the bottom is low density. The model output is in solid blue. Individual replicates of the data are in grey.

The model with our determined growth functions fit the mean proportion of females that emerged in both low and high density closely (Fig 5a). First, we fit *ν*, the relative proportion that males grow compared to females, directly and found a value of *ν* = 0.75. We then use this value to fit the growth function to the female data alone.

Although we do not fit development time based on mass, we generally saw that individuals emerge in the model around the same time as observed in the data (Fig 5b). Comparing the model to data, the large individuals in low density emerged slightly earlier. Furthermore, the modeled emergence time of the medium individuals in high density was much flatter than the individual data replicates. In all cases the model output showed lower peaks and emergence across a longer time period than found in any the graphs of individual data replicates. However, the model output was close to the average trend in the data.

### 3.4 Fixed Parameter Variation

Some of the parameters in our study were set as fixed values. We now focus on each parameter that was fixed to a particular value, and discuss its effect on the model fits when varied. The following fixed parameters arise in different variations: the resource decay rate *q*, the relative male growth *ν*, the half maximal constants (*c*_1_, *c*_2_) in the growth functions, the minimum mortality *h* in the density dependent death function, and the exponent in the density-dependent female death proportion *n*.

#### 3.4.1 Resource Decay, q

We repeated the fits for variation E using different resource decay rates, *q*. In this study, we aimed to choose a *q* that allowed all resources to be consumed in the high density treatment, but for some resources to remain in the low density treatment. We considered seven different *q* values ranging from 0.0005 to 0.05. Fig 6a left, shows how resources decay through time in our model in the low density treatment. For all values of *q* that are 0.01 or smaller, more than 20% of the original resources remained after 15 days in the low density treatment. Fig 6a right, shows resource levels through time in the high density treatment. For *q* ≥ 0.01, nearly all resources were used by day 15. Most values of *q* such that 0.005 ≤ *q* ≤ 0.2 would sufficiently fit our desired condition: complete loss of resources in high density, but not in low density. Our choice of *q* = 0.01 for this study falls within this range.

**Fig 6.**
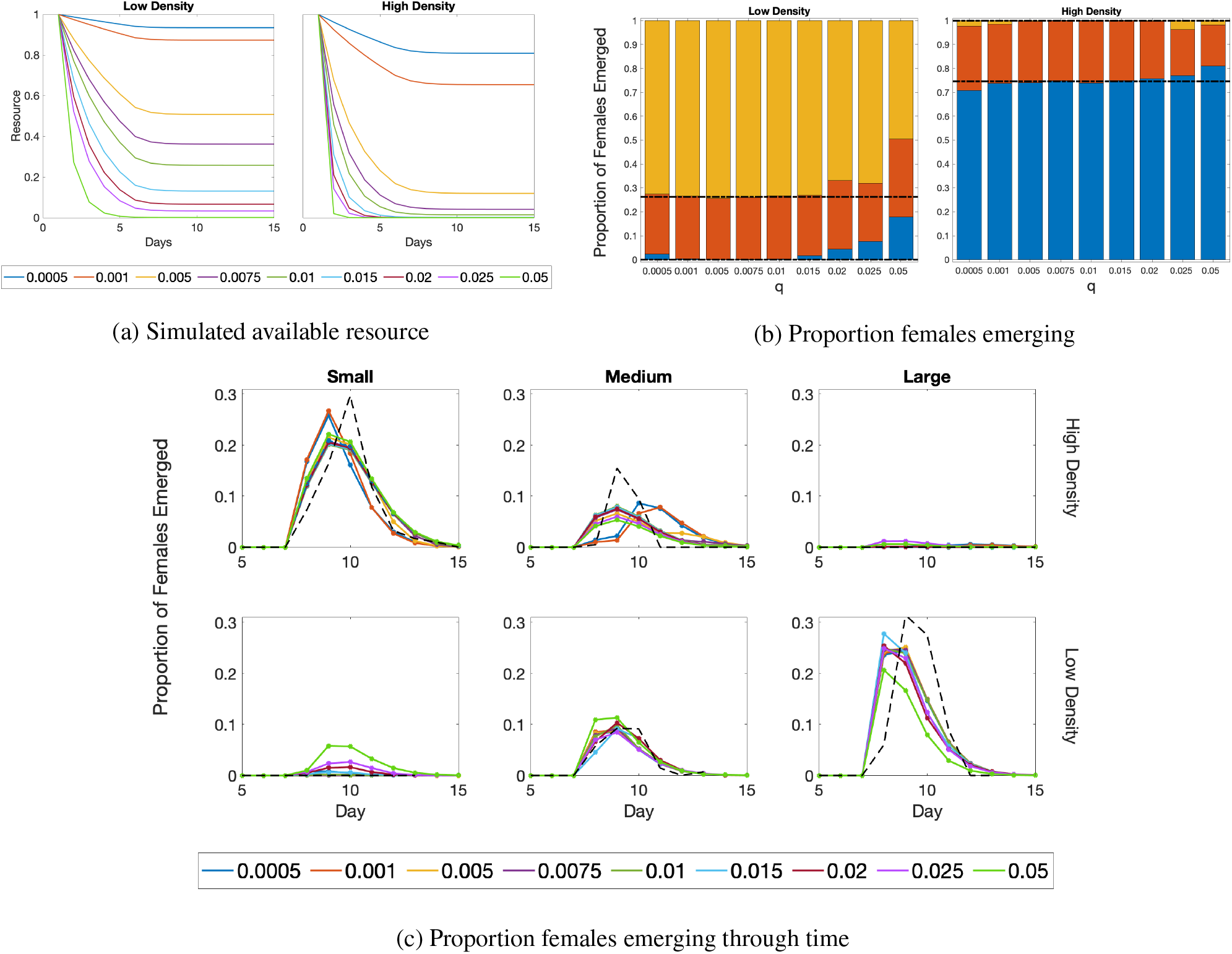
Proportion of females that emerge as resource decay rate, *q*, varies. (a) Simulated available resources throughout the time of the experiment for different values of *q*. (b) The proportion of emerging females in each mass group as *q* varies. Blue, red, and gold represent small, medium, and large mass groups, respectively. The black dashed lines indicate the divisions at which different mass groups were expected based on means of proportions of the mass groups from the data. In particular, the lower dashed line separates small and medium mosquitoes, and the upper dashed line separates medium and large mosquitoes. For close fits to the data, the blue bar would be below the lower dashed line, the red bar would be entirely between the two dashed lines, and the gold bar would be above the higher dashed line. (c) The proportion of females emerging over time by mass group: small (left), medium (middle), and large (right). The top row is the high density treatment, and the bottom row is the low density treatment. The solid color lines are model output with different *q* values. The black dashed line represents the mean of the data.

The timing of female emergence of each mass group is similar for all intermediate *q* values. For the smallest two *q* values, *q* = 0.0005 and *q* = 0.001, there were noticeable differences in timing of female emergence in the high density treatment (Fig 6c). The small mass group was slower to emerge and the medium mass group was slightly faster compared to other *q* values as well as to the average trend of the data (Fig 6c, black dashed line). For *q* = 0.05, the function did not obtain similar proportions to that seen in the data (Fig 6b). This occurred to a lesser extent for= *q* = 0.025 and *q* = 0.02. Overall, for choices of *q* that are small enough but not too small, the proportion of females emerging through time by each mass group was close to the average of the data.

#### 3.4.2 Relative Male Growth, *ν*

The value *ν* is the proportion of growth of males relative to that of females. Our model focuses on female body size and time of emergence, and *ν* does not affect the development time or growth of females. The growth of females does implicitly depend on the number of males through total larval density, but does not change regardless of the mass group of each male. In order to confirm this, we varied *ν* between 0.5 and 1, and compared female mass and total population size. The results are identical, in all aspects, for all values of *ν* in this range, except for the proportion of males in each stage (not shown).

#### 3.4.3 Per Capita Resource for Half Maximal Growth, *c*_1_ and *c*_2_

For the majority of this study, we fixed the constants of the location of half maximal growth, *c*_1_ and *c*_2_, to equal 0.5 in both growth functions (see section 2.2.4). As we want values of the growth function to approach a maximum of one as 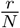 gets near one, we choose the constants of half maximal growth to occur when 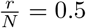. We now consider nine pairs for *c*_1_ and *c*_2_ with combinations of *c_i_* in {0.3, 0.5,0.69 ≈ log(2)}. From the combinations examined, ci = 0.5 resulted in the proportions of emerging females of small, medium, and large mass that fit the average of the data regardless of whether *c*_2_ = 0.3, 0.5 or 0.69 (S1 Fig a). The choice of *c*_1_ = 0.69 appropriately determined the proportion of emerging females in the high density treatment, but *c*_1_ = 0.69 resulted in more small mosquitoes in the low density treatment than observed in the data. We saw no difference in timing based on the choice of *c_1_* and *c_2_* (S1 Fig b).

#### 3.4.4 Minimum Mortality in Female Density-Dependent Death, *h*

We fixed the constant *h* in the numerator of the female density dependent function to be *h* = 100 (see section 2.2.3). A positive choice for *h* ensures that the death proportion is not zero at low population size. However, the final survival proportion is quite insensitive to the value for minimum mortality, h. This is because the denominator contains a large number, e.g. *f*^3^ = 24^3^. Alternative choices for *h* (up to *h* = 1000) produced very similar curves (S2 Fig). Once *h* = 2000, there were observable differences, but mostly when population levels were very low. As population levels were only low late in experiments, different values of *h* did not appreciably change the final survival proportion.

#### 3.4.5 Exponent in Density Dependent Death, *n*

In the density dependent death function for females (see section 2.2.3), we fixed the Hill exponent to three, *n* = 3. Similar survival proportions occurred for any Hill exponent greater than one (S3 Fig). However, an exponent of one deviates considerably from the sex-specific survival proportion. While higher values of the Hill exponent produced nearly identical survival proportions, the choice of a Hill exponent of three was in the region of values we considered where survival proportions were not changing with changes in the exponent. Note that for each Hill exponent chosen, while the survival proportion did not differ, other parameter estimates did change.

#### 3.4.6 Density Dependent Parameters *a, b*

In Walker et. al. [55], both parameters *a* and b in the density dependent function were fitted to a different data set with *Aedes aegypti.* We used the same values in this study for the model selection for the standard choices (*a* = 0.0043, *b* = 1.61), but vary both *a* and *b* univariately here. We considered *a* = 0.001 and 0.01 with *b* = 1.61 and then *b* = 1 and 2 with *a* = 0.0042. For each parameter, we fitted the best *k_m_* and *k_f_* for each variation C1-C3 as describe previously in section 2.4.1.

Among all combinations of *a* and *b*, the best model overall based on the AIC_*c*_ remains variation C3, where there are two compartments for both L4 and pupae. In fact, variation C3 was the best choice in all parameter combinations considered, except when *a* = 0.01 with *b* = 1.61. In this case, the best model was C2, but with a higher AIC_*c*_ than when other parameters are used. Visually, this parameter choice poorly fit the data and would not be the optimal choice (S4 Fig b).

The lowest overall AIC_*c*_ was found with the parameter set of *a* = 0.001 and *b* = 1.61 (Table 3), although it is similar to to the AIC_*c*_ of the original parameter set. The three best choices (default values *a* = 0.0043 with *b* = 1.61, *a* = 0.001 with *b* = 1.61, and *b* = 2 with *a* = 0.0043) all produced very similar results (S4 Fig a, compare solid blue line with dashed-dotted lines).

**Table 3.** AIC_*c*_ values for density dependent functions with different parameters. The original parameter choice was *a* = 0.0043 and *b* = 1.61. We then consider *a* = 0.001 and *a* = 0.01, each with *b* = 1.61. Then we consider *b* = 1 and *b* = 2, each with *a* = 0.0043.

| Variation | Original | $a = 0.001$ | $a = 0.01$ | $b = 1$ | $b = 2$ |
| --- | --- | --- | --- | --- | --- |
| C1 | 22.9 | 23.1 | 23.1 | 22.9 | 23.0 |
| C2 | 22.3 | 22.4 | 22.8 | 22.4 | 22.3 |
| C3 | 21.3 | 21.1 | 23.1 | 21.8 | 21.2 |

## 4 Discussion

In this work, we developed a discrete time mathematical model parameterized with laboratory data that demonstrated how density in the larval environment affects variability in mosquito body size. Our model separates masses into three groups – small, medium, and large – and tracks mosquito growth through aquatic stages. Using our model, we determined the distribution of mass and sex at adult emergence under different larval density treatments, and we illustrated the interactions between larval environmental conditions and adult body mass, which could have important implications for mosquito population and mosquito-borne pathogen control. This work is an important contribution towards understanding how body mass affects mosquito development and how conditions in the early developmental stages may have longer-term consequences.

Larvae require more than a single day in each developmental stage. When we used only a single compartment for each aquatic stage, more than a quarter of the individuals in the model emerged earlier than the timing observed in the data. Furthermore, the model showed emergence that is distributed over many days with much lower numbers per day than seen in the data. Comparing our model variations, the AICc selected the model with a second compartment in both L4 and pupae. This model variation extends the minimum time of emergence by two days. Importantly, while forcing the minimum to be larger by two days resulted in model output with time of emergence closer to that of the data, it also produced variance that was more similar to the data. The additional time spent in L4 was suggested previously by Levi et al. [57]. In their study, they found that in nutrient rich environments larvae initially grew quickly, but then spent a longer time in L4. In less rich nutrient environments, individuals developed more slowly throughout all stages. In both situations, they noticed similar overall times until emergence. In studies using similar temperatures to our own, pupae had an average development time close to two days [58–60]. While in our model we set the minimum development time of pupae to two days, the overall development time is often longer because individuals may remain in any stage for a longer period.

We showed that including density-dependent death is important to accurately represent the development of female larvae. Our data showed that the survival of female mosquitoes is diminished in the high density treatment. Compared to the low density treatment, the difference in survival was significant enough that the AICc selected this even though the model requires two additional parameters to incorporate this density dependence. In contrast, the difference between male and female survival in our experiment was negligible compared to the difference in female survival in the two density treatments. This is emphasized further by the lower AICc score for the model with a single constant parameter representing both male and female survival compared to the model with two parameters for sex-dependent survival. Density plays a key role in females’ survival, and survival to adulthood is an important factor in mosquito population and disease dynamics. In particular, because female mosquitoes transmit pathogens, any significant alterations of female survival could propagate through population dynamic processes to cause profound impacts on population magnitude and potentially even pathogen transmission.

Our model fitting demonstrates that growth to the largest mass group quickly becomes restricted in resource-limited environments. We determined the exponent of the function determining growth from medium to large mass groups (*G*_2_) to be *n*_2_ = 9. Given that the exponent determines the steepness of the growth curve, the proportion growing from medium to large mosquitoes rapidly diminishes to nearly zero as resources decay. This indicates that only in environments with ample resources will individuals grow into the largest mass groups. It should be noted that we fixed rather than fit several parameters in part due to the lack of variation in feeding regimes in our data. To further explore the consequences of this choice, in section 3.4 we considered the sensitivity of our model to these parameters. We showed that our choices of parameters either give the best fits overall or the results were not sensitive to the parameter. While examining various feeding regimes will be important future work, our model and choices of parameters described our data well.

In the model, development time was less important in determining adult size at emergence compared to the larval density environment. Our experiments show significant differences in the emergence of different mass groups between the high and low density treatments. In particular, no large mass individuals emerged in high density and almost no small mass individuals emerged in low density. The only significant difference observed for development time as a function of mass occurred from small to medium mass groups in the high density treatment, but the effect size was small. Aznar e.t. al. [30] modeled the importance of reaching a particular size before pupation could occur. However, they did not consider how size at emergence varied, and commented that the variability in size was incidental and based more on time to emergence. In a modeling investigation, Gilpin and McClelland considered a uniform distribution for size, and found the range of the spread in size increased linearly over time [31]. This quickly led to very large and very small mosquitoes with equal probability, and there was no relationship between size and larval environment. In other work, Padmanabha et. al. aimed to predict time of pupation, but not variation in mass at emergence [32]. They did consider variation in time to emergence as well as survival by temperature, but did not track changes in mass. In contrast, our model focuses on the distributions of mosquito mass with different larval environments.

Our model is limited in two key ways: we focus on a single mosquito species, *Aedes aegypti*, and we assume a constant temperature setting. While there are other species of the genus *Aedes* that typically have similar behavior, even within the *Aedes* genus there are differences that could change the results. Extrapolations to other species would require experiments for species-specific parameterization. Temperature has a key role in development time and mortality [58,59], and inclusion of temperature variation would improve the model’s utility. Adding in effects of temperature would significantly increase the complexity of the model; however, it will be important for future iterations of the model to consider variation in temperature. In addition, features found to be important in other models have been omitted because we focus on results at emergence rather than specific results at each stage. For example, the inclusion of resource dependent mortality at young stages and resource dependent delay of L4 would more accurately describe behavior of individual stages. Additionally, we use mass as a measurement in our model, while Padmanabha et. al. [32] found that reserves, rather than raw weight in the model, more accurately describe when individual mosquitoes pupate. While reserves are an important indicator of success as an adult as well, using mass as a proxy still performs well and is easier to measure.

The work described herein is an important contribution towards understanding how environmental conditions during juvenile growth affect mosquito mass and development, and thus control of mosquito populations and the diseases whose causative agents they transmit. Our model can be adapted to consider the importance of larval environmental heterogeneity on mosquito mitigation strategies. Many traditional and novel control methods focus on reductions in the larval population or late-acting lethal measures that allow mosquitoes to complete the majority of the juvenile stage before death [61–64]. Our work suggests that these control approaches could have varying success as disease mitigation strategies if larval dynamics are altered in such a way that increases, for example, female body size or overall survival. Although we did not model how mass affects the adult population dynamics or disease spread, our work modeling mass-dependent aquatic development of mosquitoes is useful for studying mosquito mitigation strategies regardless of the life stage which they directly affect. For example, if a control method targets adults only, then adult mosquito population drops. A lower adult population leads to fewer eggs laid, which in turn leads to lower aquatic density. Given lower density environments, mosquitoes that emerge may tend to be larger, and the females will have better survival [48]. Thus, adult population reduction may not lead to the intended consequences and could in fact have an adverse effect on mosquito control efforts. Given this, it is critical to better understand how mass at emergence is determined by larval environments and to expand on this to increase our understanding of the robustness of mosquitoes as potential vectors for disease spread. Furthermore, this emphasizes that integrated control approaches that target multiple life stages are important because mitigation strategies focusing on a single life stage may have a number of shortcomings.

## 5 Supporting information

**S1 Fig.**
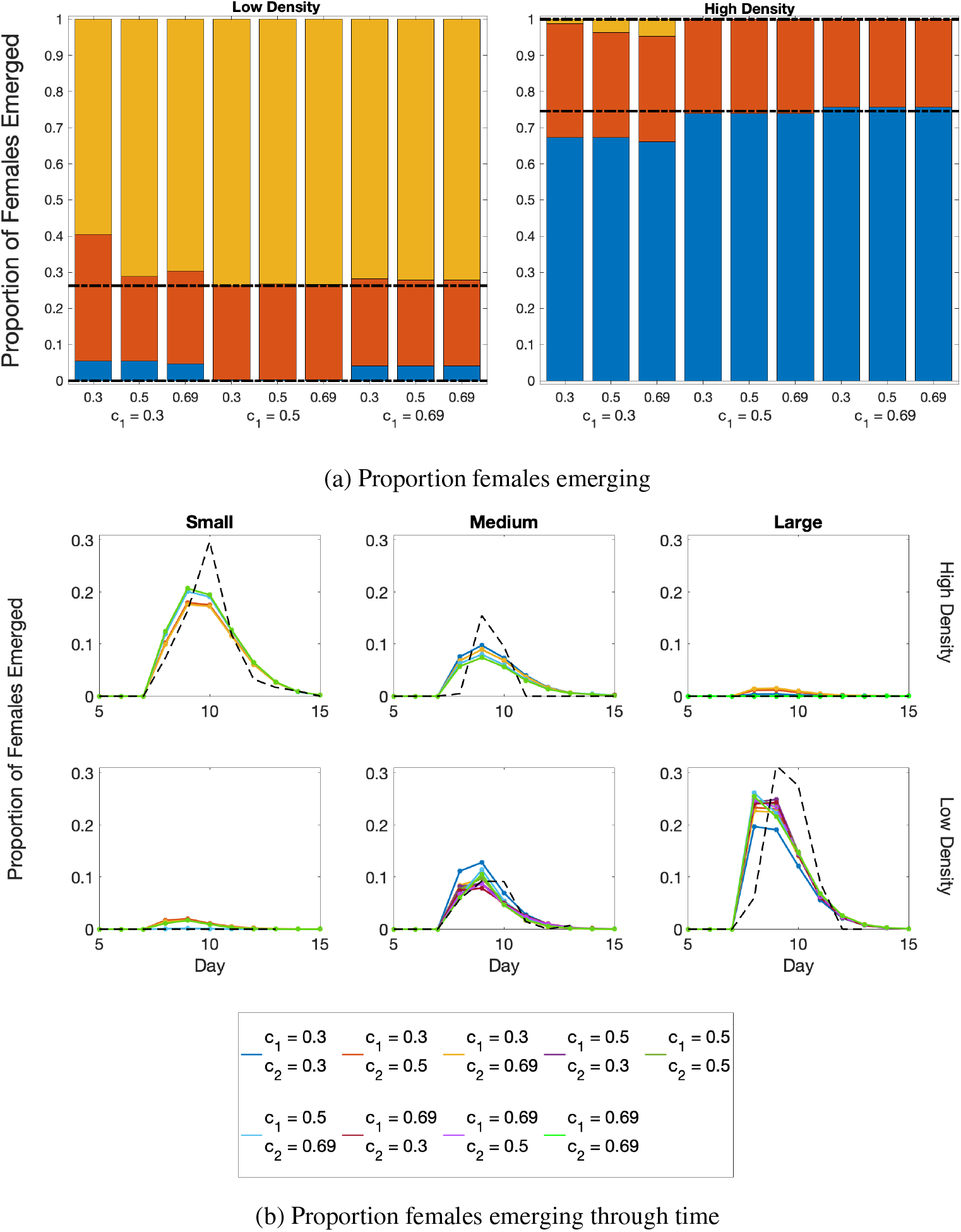
Proportion of females that emerge as half maximal growth varies. (a) The proportion of emerging females in each mass group as *c*_1_ and *c*_2_ vary. Blue, red, and gold represent small, medium, and large mass groups, respectively. The black dashed lines indicate the divisions at which different mass groups were expected based on means of proportions of the mass groups from the data. In particular, the lower dashed line separates small and medium mosquitoes, and the upper dashed line separates medium and large mosquitoes. For close fits to the data, the blue bar would be below the lower dashed line, the red bar would be entirely between the two dashed lines, and the gold bar would be above the higher dashed line. (b) The proportion of females emerging over time by mass group: small (left), medium (middle), and large (right). The top row is the high density treatment, and the bottom row is the low density treatment. The solid color lines are model output with different *c*_1_ and *c*_2_ values. The black dashed line represents the mean of the data.

**S2 Fig.**
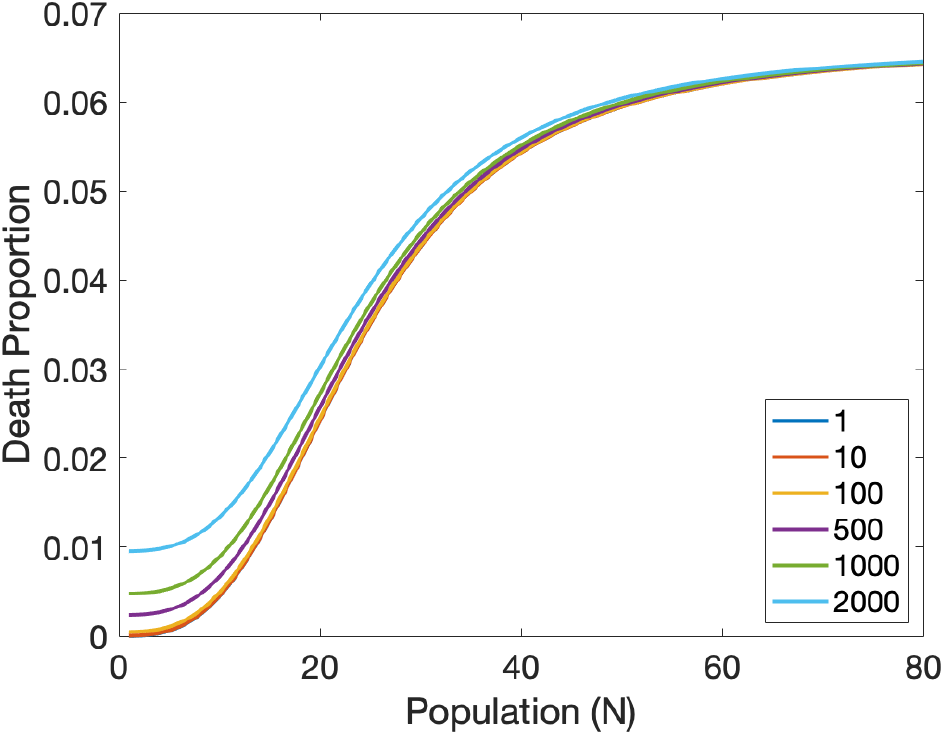
Density-dependent death proportion as the minimum death varies. Density dependent death function 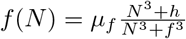 with *f* = 24 and *μ_f_* = 0.066. The minimum constant *h* varies from 1 to 2000. See Section 2.2.3 for details on the functional form.

**S3 Fig.**
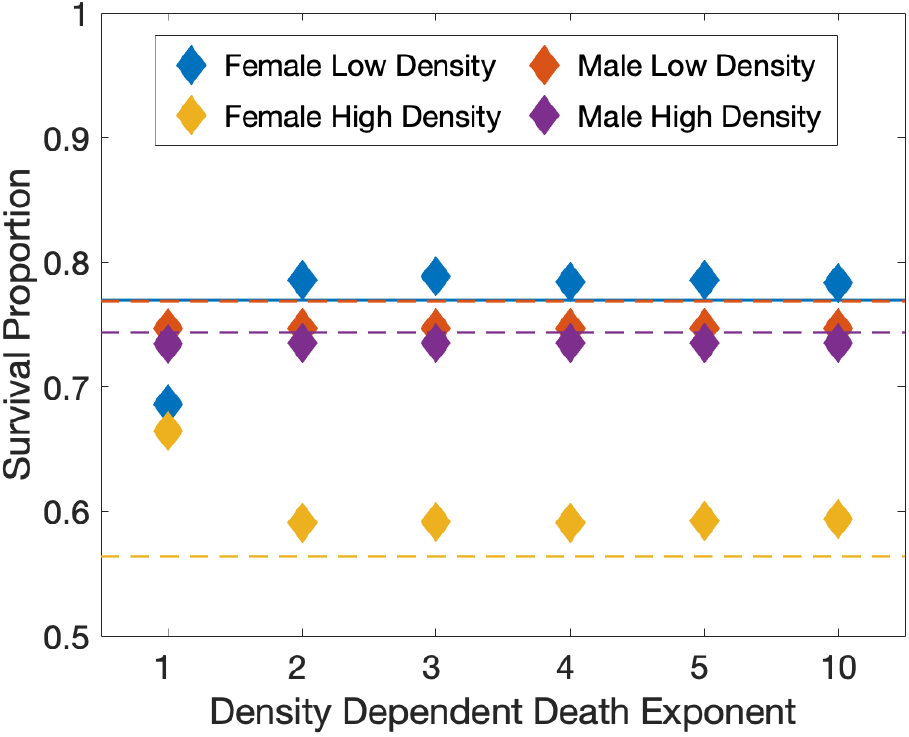
Density-dependent death proportion as the Hill exponent changes. Model results employing the density dependent death function 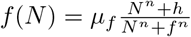 with *f* = 24, *μ_f_* = 0.066, and *h* = 100. The density dependent death exponent, *n*, varies along the x-axis from 1 to 10. The total larvae at a given time, *N*, changes in the course of the model simulations. The dashed lines represent the median values from the data and the diamonds the model results for the survival proportion of females in low density (blue), females in high density (yellow), males in low density (red), and males in high density (purple). The survival proportion for males and females in low density is indistinguishable in the data. See section 2.2.3 for details on the functional form of *f*(*N*).

**S4 Fig.**
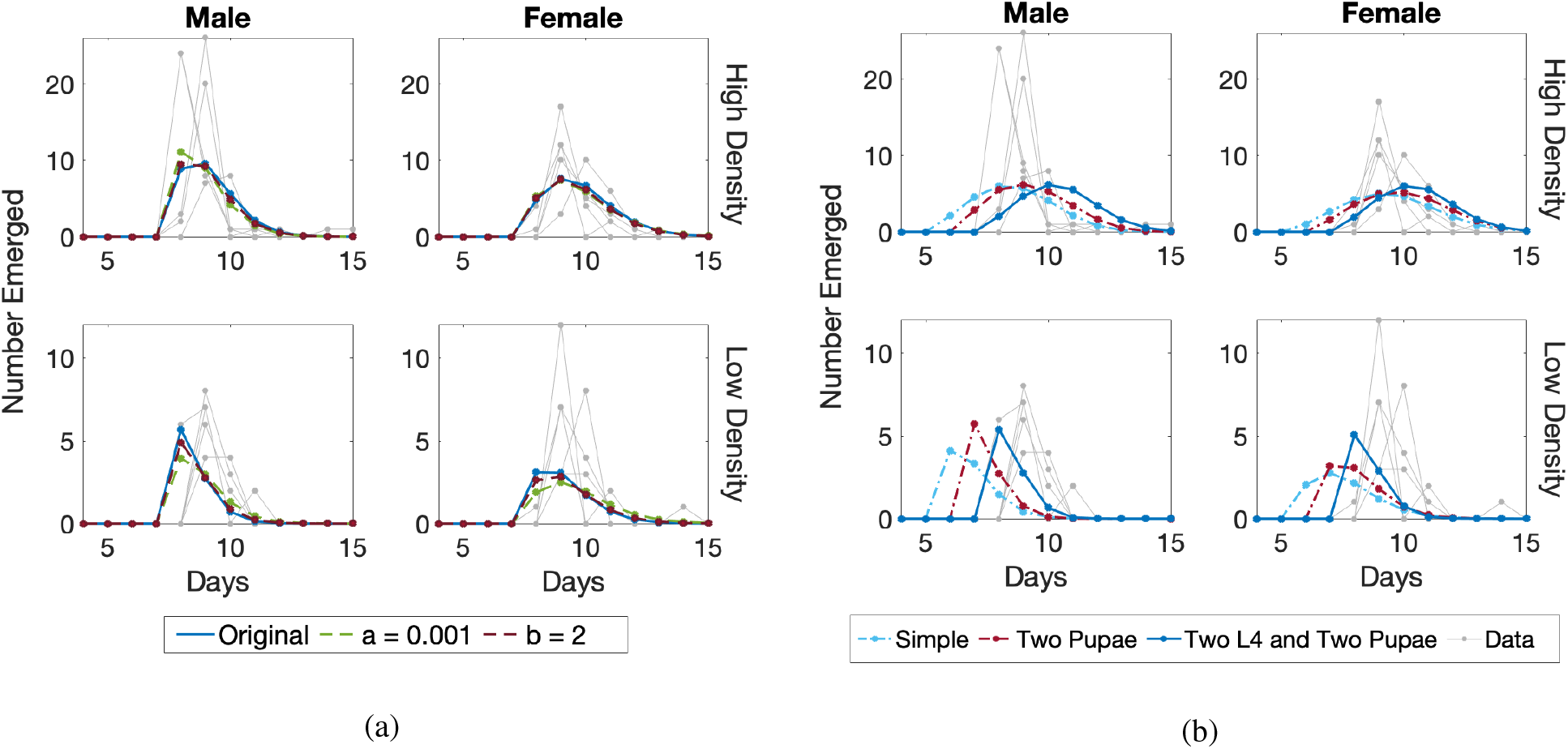
Varying parameters *a, b* in the density dependent function. (a) The solid blue line shows the choice of our model with our original parameters *a* = 0.0043 and *b* = 1.61. The two other parameter choices with similar AIC_*c*_ values are shown, when *a* = 0.001 and *b* = 1.61 (dashed green line) and when *a* = 0. 0043 and *b* = 2 (dashed dark maroon line). (b) This shows all three variations C1 (dashed dotted light blue line), C2 (dashed dotted maroon line), and C3 (solid blue line) with the parameters set at *a* = 0.01 and *b* = 1.61.

## 6 Acknowledgments

CV was supported by the USDA, National Institute of Food and Agriculture, Hatch project 1017860. CV, KC, LC, and MW received support from the Center for Emerging, Zoonotic, and Arthropod-borne Pathogens at Virginia Tech. LC and MW were supported by NSF Grant 1853495

